# Molecular and Functional Analysis of Calcium Binding by a Cancer-linked Calreticulin Mutant

**DOI:** 10.1101/2025.06.25.661618

**Authors:** Ishmael Nii Ayibontey Tagoe, Amanpreet Kaur, Osbourne Quaye, Emmanuel Ayitey Tagoe, Nicole Koropatkin, Leslie S. Satin, Malini Raghavan

## Abstract

Calreticulin (CRT) is an endoplasmic reticulum (ER) chaperone with low affinity calcium binding sites in its C-terminal domain. This region is altered by somatic mutations in the CRT gene (*CALR*), which drive a subset of myeloproliferative neoplasms (MPN). Perturbations in ER calcium storage and signaling are reported for the MPN type I mutant, CRT_Del52_, and are linked to disease pathogenesis. Using recombinant CRT proteins, we found similar low affinity calcium binding characteristics for wild-type CRT and CRT_Del52_, as determined by isothermal titration calorimetry (ITC). Residues 340–349, conserved in both wild-type CRT and CRT_Del52_, contribute to binding. Furthermore, ER and cytosolic calcium levels and store-operated calcium entry (SOCE) were comparable in CRT knockout (CRT-KO) cells reconstituted with wild-type CRT or CRT_Del52_. Notably, the CRT-KO induces expression of multiple ER proteins known to contain low affinity calcium binding sites or regulate ER calcium levels. Overall, these findings indicate that CRT_Del52_ retains at least some low affinity calcium binding capacity and that CRT-deficiency induces compensatory cellular changes that maintain ER calcium homeostasis.

**Summary:** Pathogenic CRT_Del52_ retains at least partial low-affinity calcium-binding capacity. Additionally, live-cell imaging and flow cytometry with ratiometric calcium probes indicate that ER and cytosolic calcium levels, as well as store-operated calcium entry (SOCE), are comparable in CRT-deficient cells rescued with either wild-type CRT or CRT_Del52._

## Introduction

Calreticulin (CRT) is an endoplasmic reticulum (ER) luminal protein that plays key roles in the assembly and folding of N-linked glycoproteins. CRT has a single high affinity calcium binding site within a globular domain and a highly acidic C-terminal domain that contains multiple low affinity calcium binding sites (Baksh and Michalak, 1991, Wijeyesakere et al., 2011). As an ER chaperone, CRT is thought to play a critical role in maintaining cellular calcium homeostasis (Venkatesan et al., 2021b, Michalak, 2024). The ER has a relatively high free calcium concentration of 0.5-1 μM through the presence of various low affinity calcium-binding proteins that buffer calcium, whereas cytosolic calcium levels are maintained within a narrow range of around 100 nM (Michalak, 2024). The maintenance of calcium homeostasis within cellular compartments is essential for cell function and survival (Bagur and Hajnoczky, 2017). Early studies have shown that the C-domain of CRT contributes to ER calcium storage, although free ER luminal concentrations are unchanged (Nakamura et al., 2001).

Somatic mutations of exon 9 of the *CALR* gene including a 52-base pair deletion (Del52; type I mutation) and a 5-base pair insertion (Ins5; type 2 mutation), are driver mutations in some myeloproliferative neoplasms (MPN), specifically in essential thrombocythemia (ET) and primary and secondary myelofibrosis (PMF) (Klampfl et al., 2013, Nangalia et al., 2013). The MPN mutations change the CRT C-domain residues from being predominantly acidic to a high prevalence of basic amino acids and result in a loss of the ER-retention KDEL sequence (Klampfl et al., 2013, Nangalia et al., 2013) (Figure 1). Although all MPN CRT mutants change the sequence and charge of the C-domain, a greater number of acidic residues are retained in MPN type 2 mutant (CRT_Ins5_), compared to the type I mutant (CRT_Del52_) (Figure 1). The loss of more acidic residues from the C-terminal domain of CRT_Del52_ has been reported to impair its calcium binding and reduce ER calcium levels (Ibarra et al., 2022). Store-operated calcium entry (SOCE) is a physiological process that allows for the entry of extracellular calcium and ER calcium restoration following ER calcium depletion (Putney, 2011, Zhang et al., 2020). Altered calcium release from the ER and SOCE was reported for MPN type I but not type 2 mutants in cultured megakaryocytes from MPN patients (Pietra et al., 2016). In another study, spontaneous calcium spikes and increased SOCE activation were observed in cultured megakaryocytes from MPN type I and type 2 patients when compared to healthy donors (Di Buduo et al., 2020).

**Figure 1:**
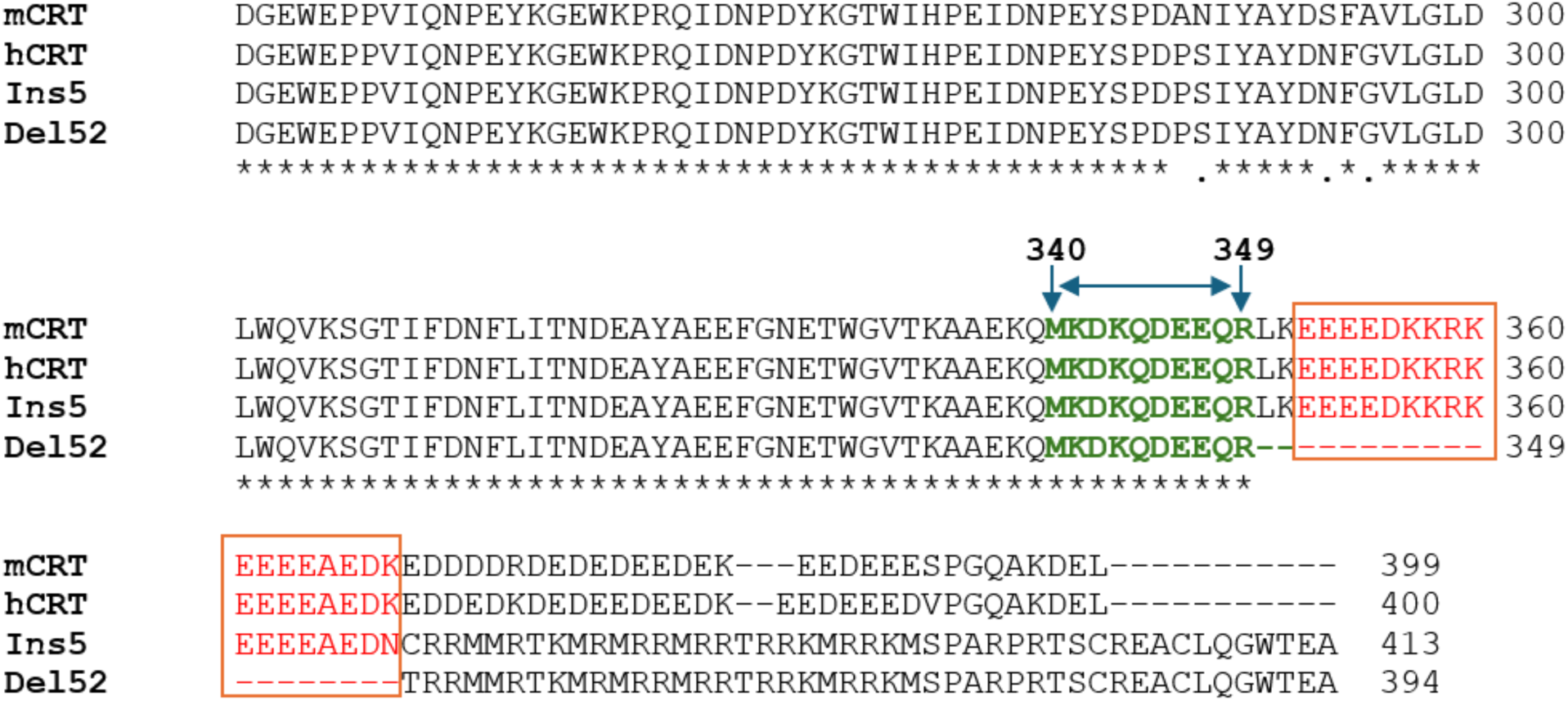
Sequence alignments of the C-terminal residues of murine and human calreticulin (mCRT and hCRT), and the MPN-linked CRT mutants Ins5 and Del52. Multiple sequence alignment using Clustal Omega was employed to compare C-domain sequences of mature mCRT, hCRT, Del52, and Ins5 proteins. Acidic residues (340 – 349) previously implicated in low affinity calcium binding by mCRT that are shared between wild-type and both CRT_Del52_ and CRT_Ins5_ are highlighted in green. Additional acidic residues shared just between wild-type CRT and CRT_Ins5_ are shown in red. There are additional acidic residues unique to wild-type CRT. Mature protein numbering is used for the CRT sequences.

Although global calcium binding defects were predicted or qualitatively measured for CRT_Del52_ (Shivarov et al., 2014, Ibarra et al., 2022) and altered cellular calcium signaling is also reported (Pietra et al., 2016, Di Buduo et al., 2020), it remains unclear whether MPN-linked CRT_Del52_ has any low affinity calcium binding capacity, given that a few conserved acidic residues are retained in its C-domain (Figure 1). In fact, our previous studies showed that a truncated version of murine wild-type CRT (mCRT) containing the non-mutated part of CRT_Del52_ and two additional C-terminal residues (mCRT_1-351_) showed similar low affinity calcium binding characteristics as the wild-type mCRT (Wijeyesakere et al., 2011). On the other hand, further truncation of mCRT (mCRT_1-339_) was needed to disrupt calcium binding (Wijeyesakere et al., 2011, Wijeyesakere et al., 2016). The goal of the current study was to quantitatively measure low affinity calcium binding by CRT_Del52_ compared with wild-type human CRT and also quantify ER and cytosolic calcium levels and calcium responsiveness in cells expressing wild-type CRT or CRT_Del52_.

## Results

### MPN-linked CRT_Del52_ retained at least some low affinity calcium binding sites

Previous studies (Wijeyesakere et al., 2011) have implicated residues 340 to 351 of murine CRT, which are identical in wild-type human CRT and fully conserved with the 340 to 349 residues of CRT_Del52_, to be involved in low affinity calcium binding. Further truncation of the C-domain to the residue 339 (mCRT_1-339_) led to the loss of calcium binding measured by ITC (Wijeyesakere et al., 2011). Figure 1 shows a sequence alignment of murine CRT, human CRT, CRT_Ins5,_ and CRT_Del52_ proteins and indicates that the residues previously implicated in low affinity calcium binding are also shared between wild-type CRT and CRT_Del52_. Recombinant his and GB1 tagged purified proteins were used to compare calcium binding by human CRT and its variants. Chromatograms for human wild-type CRT, CRT_Del52_, CRT_(1-339)_ and CRT_(1-351)_ are shown in Figure 2A. SDS-PAGE gels corresponding to the purified proteins are shown in Figure 2B. Intact protein mass spectrometry for CRT_Del52_ indicated 35-42 amino acid truncations relative to the full protein sequence for different preparations, possibly from C-terminal protein degradation in *E. coli*. The truncated proteins still include acidic residues (340–351) previously implicated in low affinity calcium binding by murine CRT that are shared between wild-type and CRT_Del52_ (Figure 1).

**Figure 2:**
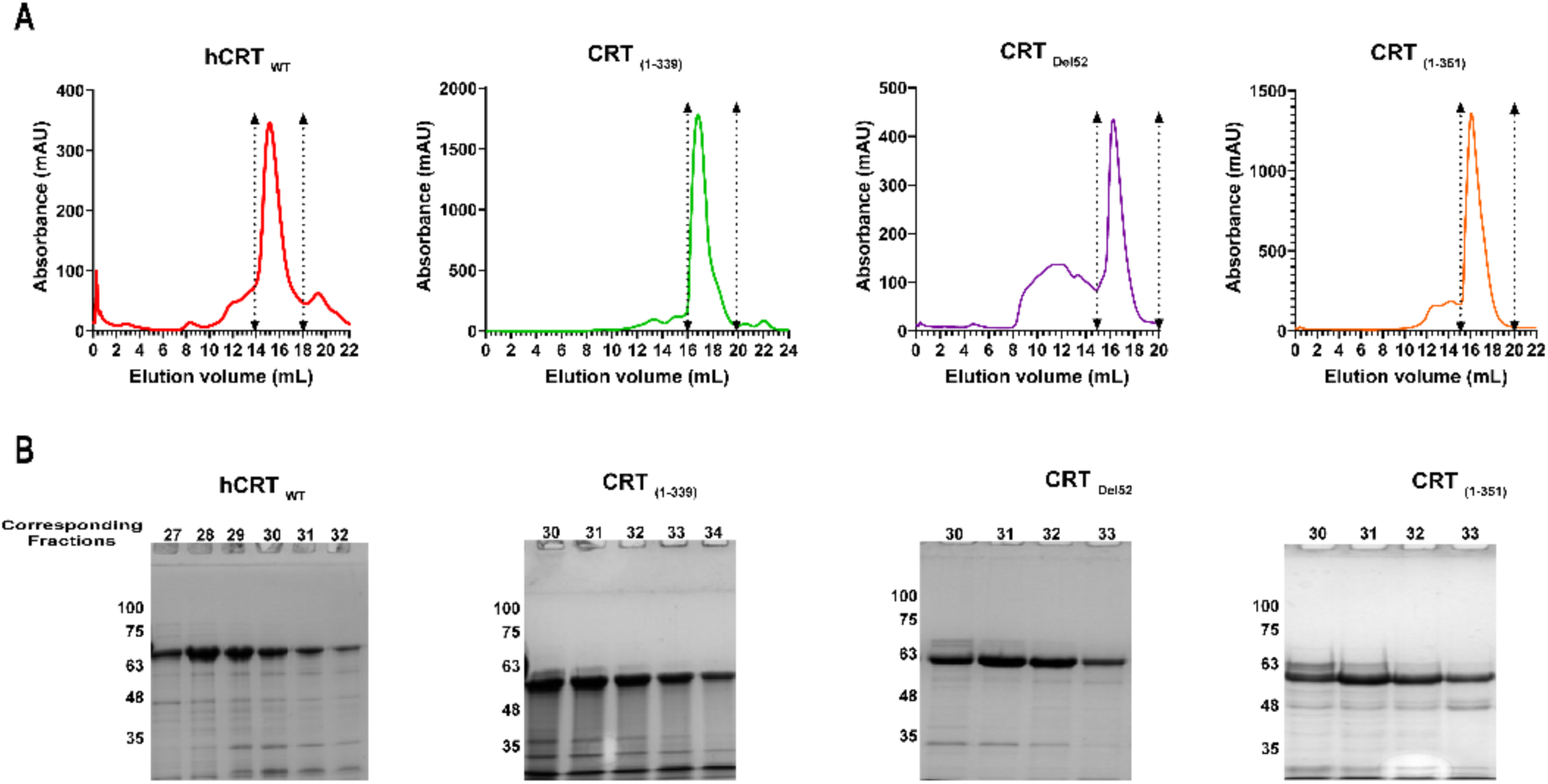
Purification of recombinant human wild-type CRT, CRT_Del52,_ and C-domain truncation mutants. **A)** Representative chromatograms of recombinant GB1-and his-tagged human CRT_WT_ (hCRT_WT_), CRT_Del52_, and the C-domain truncation mutants hCRT _(1-351)_ and hCRT _(1-339)_ purified by gel exclusion chromatography after expression in *E. coli* and initial purification using a nickel resin. **B)** Representative Coomassie blue-stained SDS-PAGE gels of fractions corresponding to the monomer peak (those contained within the region indicated by the double headed arrows on the chromatogram) of purified proteins that were used for calcium-binding measurements.

CRT_Ins5_ was not purified in sufficient quantities needed for ITC experiments and was not further investigated in this study.

For the ITC measurements, calcium titrations were performed after pre-blocking the high affinity calcium binding sites with 50-100 µM CaCl_2_. In a previous study (Wijeyesakere et al., 2011), we showed that at a starting calcium concentration of 0 mM and with CaCl_2_ injections to a final concentration of 70-80 μM, the measured K_D_ value was 16.6 μM for calcium binding to wild type murine CRT, (which has ∼95% sequence identity with human CRT), corresponding to the high affinity site. On the other hand, at a starting calcium concentration of 50 μM and CaCl_2_ injections (at 33 μM each) to a final concentration of 700-850 μM, the measured K_D_ value for calcium binding to wild type murine CRT was 590 μM (corresponding to the low affinity sites). We did not measure the high affinity sites under the latter conditions (Wijeyesakere et al., 2011). In the present experiments, at starting CaCl_2_ concentrations of 50-100 μM and 25 sequential 10µL injections of 76 μM CaCl_2_ to a final concentration of 1650-1700 μM, similar binding profiles and K_D_ values in the 400-1000 μM range were measured with wild-type human CRT, CRT_Del52,_ and the CRT_(1-351)_ mutant (Figures 3A, C and D and Table 1). However, consistent with previous studies with mCRT_(1–339)_, endothermic calcium binding by human CRT_(1–339)_ was of lower magnitude compared to other constructs, with a K_D_ value that was too low to be reliably fit, suggesting the loss of low affinity calcium binding sites in this mutant (Figure 3B and Table 1). There are exothermic peaks observed with human CRT_(1–339)_ at the beginning of the calcium titration, which may be attributed to altered low affinity sites retained within the mutant or reduced affinity of the high affinity sites. We concluded that CRT_Del52_ retains at least some low affinity calcium binding sites that are measurable by ITC.

**Figure 3:**
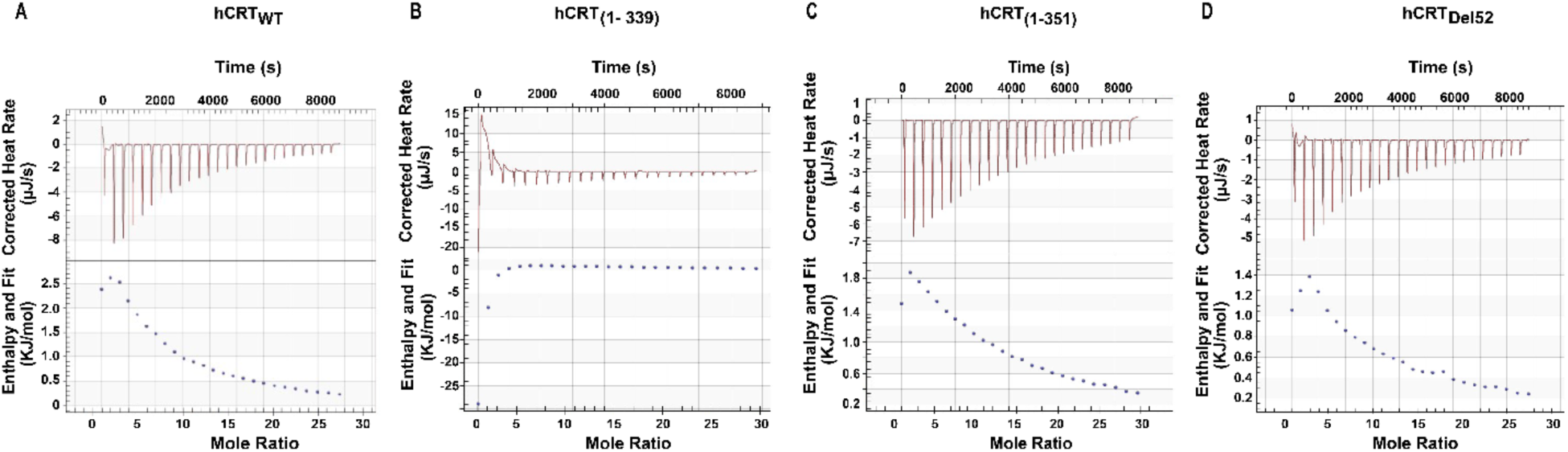
Calcium binding to the low affinity sites of purified human CRT proteins. Low affinity calcium binding by purified hCRT_WT_, CRT_Del52_, and the indicated truncation mutants was measured using isothermal titration calorimetry. **A-D)** Representative isotherm and fitting curves of calcium binding to the low affinity sites of the wild-type and mutant CRT proteins as indicated, measured using isothermal titration calorimetry (ITC). Prior to titration, the proteins were incubated with 50-100 µM CaCl_2_ to block the high affinity calcium binding sites. At 37°C, 10 mM CaCl_2_ was titrated against 100 µM protein. The heat change was measured for 25 injections, and the data were curve-fitted to calculate the dissociation constant, enthalpy, and binding stoichiometry using NanoAnalyze software. Data replicates and statistical analyses are specified in Table 1.

**Table 1:** Calorimetric data on measurement of calcium binding to the low affinity binding site of recombinant purified CRT.

| Recombinant CRT proteins/Isothermal calorimetric measurement | $\Delta H$ (kJ/mol) | n (mol) | Dissociation constant - $K_D$ ($\mu M$) |
| --- | --- | --- | --- |
| CRT<sub>WT</sub> | $7.88 \pm 2.15$ | $5.40 \pm 1.31$ | $574.24 \pm 109.93$ |
| CRT<sub>(1-351)</sub> | $79.82 \pm 32.55$ | $2.59 \pm 1.56$ | $444.34 \pm 139.99$ |
| CRT<sub>(1-339)</sub> |  |  | Too low to reliably fit |
| CRT<sub>Del52</sub> | $3.89 \pm 1.36$ | $7.73 \pm 1.81$ | $938.82 \pm 275.89$ |

Calorimetric data are represented as the mean ± S.E. Data shown are averages from 3 or 5 independent replicates.

### Similar ER calcium levels in CRT-KO HEK cells reconstituted with wild-type CRT or CRT_Del52_

While the CRT_Del52_ mutant retains some ability to bind calcium (Figure 3 and Table 1), the loss of the ER-retention KDEL sequence induced by the frameshift mutation, which results in the secretion of the protein from the ER (Araki et al., 2016, Chachoua et al., 2016, Han et al., 2016, Arshad and Cresswell, 2018, Liu et al., 2020, Masubuchi et al., 2020, Venkatesan et al., 2021a, Pecquet et al., 2023) might be expected to cause ER calcium insufficiency in cells expressing CRT_Del52_ compared to wild-type CRT. We investigated ER calcium signals in CRT knock-out HEK293T cells generated by CRISPR/Cas9 gene editing. A clonal KO line was generated and subsequently reconstituted with constructs encoding wild-type CRT, CRT_Del52_, or CRT_Del52_ with a KDEL sequence added at the C-terminus (CRT_Del52-KDEL_). Protein expression was assessed by immunoblotting with anti-CRT(N), which detects an N-domain epitope present in all CRT constructs, and anti-CRT(C-mut), which was specifically raised against the mutant CRT C-terminus (Venkatesan et al., 2021a). While the recognition with anti-CRT (N) was very weak in lysates of cells expressing CRT_Del52,_ which is secreted, CRT_Del52-KDEL_ is readily detectable with both antibodies, consistent with prior studies demonstrating the contributions of the KDEL sequence to the ER retention of CRT and its mutants, including in HEK cells (Arshad and Cresswell, 2018, Sonnichsen et al., 1994). Both wild-type CRT and CRT_Del52-KDEL_ were reconstituted to higher levels than in the parental cells (Figure 4A, lanes 1 and 2 compared to 7-8 and 11-12).

**Figure 4:**
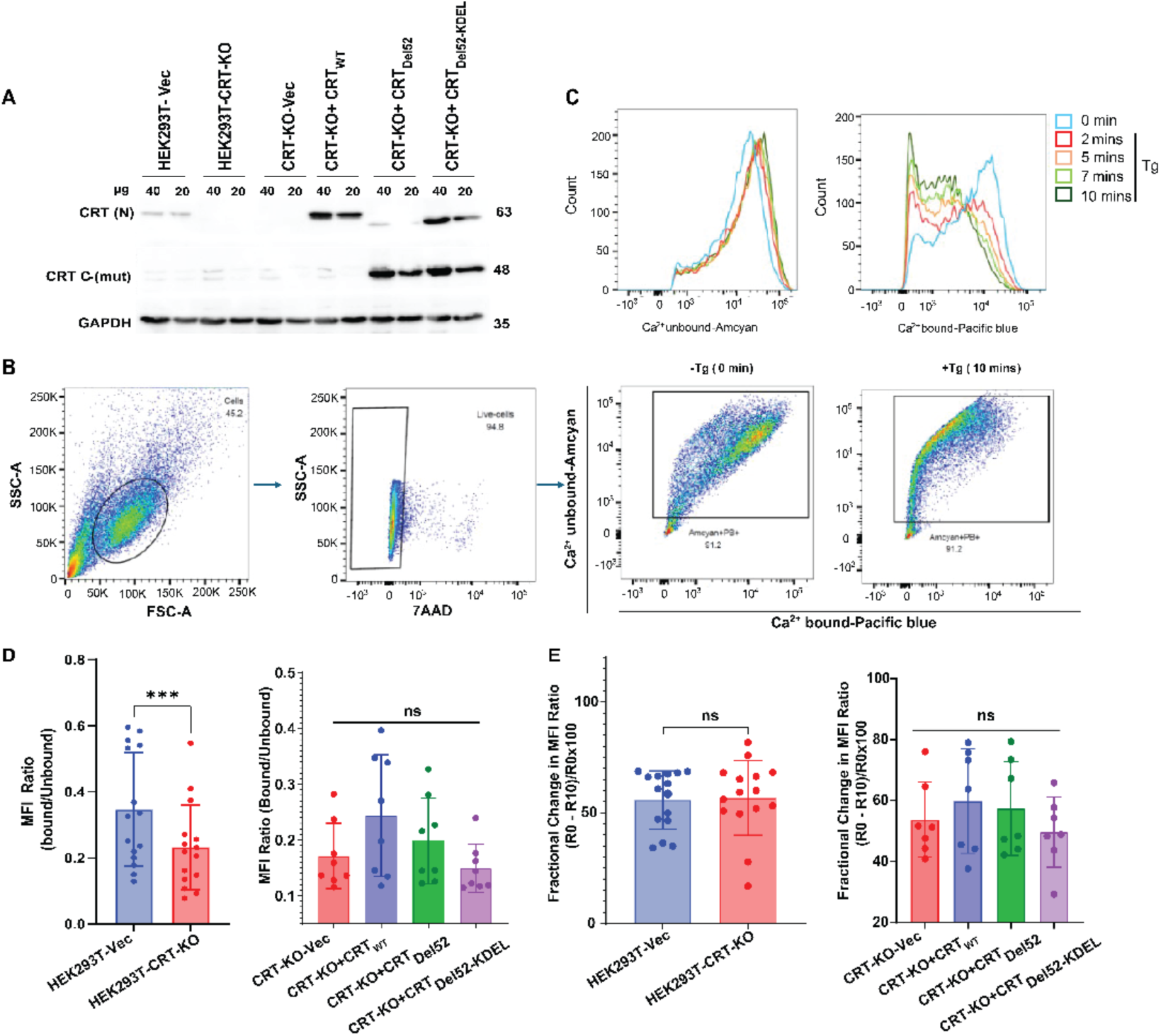
Flow cytometry-based ER calcium measurements indicate similar ER calcium levels in CRT-KO HEK293T cells and those reconstituted with wild-type CRT and CRT_Del52_. **A)** CRISPR-Cas9 editing of HEK cells using lentiviral constructs with calreticulin-targeting gRNA (CRT-KO) or empty vector (HEK-Vec). CRT-KO HEK cells were subsequently reconstituted with wild-type human CRT, CRT_Del52,_ or CRT_Del52_ with a C-terminal KDEL sequence as indicated. Representative blots showing expression levels of CRT in the indicated HEK cells. Cell lysates were subjected to SDS-PAGE followed by immunoblotting. Wild-type CRT, mutant CRT, and GAPDH were detected with anti-CRT (N), anti-CRT C-(mut), and anti-GAPDH antibodies, respectively. **B)** Gating strategy for flow cytometry. HEK-Vec cells transiently transfected with the vector encoding the ratiometric GEM-CEPIA1er probe for 72 hours were used to assess ER calcium levels by flow cytometry based on the measurements of fluorescence emission signals for the Ca^2+^-bound or unbound probe in the Pacific Blue and Amcyan channels, respectively following excitation with the 405 nm (violet) laser. Transfected cells were gated based on forward (FSC-A) and side scatter (SSC-A), then on 7AAD-negative (live) cells, and further on the cells positive for the calcium-bound and unbound probe signals. Representative dot plots of changes in fluorescence intensities in response to treatment with thapsigargin (Tg), a SERCA inhibitor, before (0 mins) or after (10 mins) treatment, are shown. **C)** Representative histograms showing changes to Ca^2+^-bound and unbound signals at different time points following Tg treatment of cells. Mean fluorescence intensities (MFI) for Ca^2+^-bound and unbound signals were used to calculate the bound/unbound ratios at different time points. **D and E)** The basal free ER calcium signals were measured as bound/unbound MFIs ratios from the corresponding histograms of CRT-KO or HEK-Vec cells (D, left panel) or CRT-KO cells reconstituted with the indicated CRT constructs (D, right panel). The fractional changes in ER calcium signals in response to Tg were measured in the indicated cells (E). The fractional change ratios were calculated as (R_0_-R_10_/R_0_)x100. Statistical significance was determined using GraphPad Prism and based on paired t-tests (left panels of D and E) or repeated measures one-way ANOVA analyses with Tukey’s test (right panels of D and E). ns, not significant; *** P value < 0.001. All measurements were undertaken in the presence of 2 mM extracellular Ca^2+^. Data shown in D and E are based on 7-15 independent experiments as indicated by the individual data points within the graphs. GAPDH, glyceraldehyde-3-phosphate dehydrogenase; SDS-PAGE, sodium dodecyl sulfate-polyacrylamide gel electrophoresis.

The KO and reconstituted cells were further transfected with pCIS-GEM-CEPIA1er, which encodes a ratiometric ER calcium probe (Suzuki et al., 2014). Emission signals for calcium-bound (peak ∼ 460 nm) and unbound (peak ∼ 510 nm) probe were measured using flow cytometry in the Pacific Blue and AmCyan channels, respectively, following excitation with the 405 nm (violet) laser. Flow cytometric analysis was undertaken in a buffer containing 2 mM extracellular Ca^2+^ (Figures 4B and 4C). There was a significant decrease in basal free ER calcium, measured by the bound/unbound ratios in CRT-KO cells compared to the parental cells, (Figure 4D, left panel). ER calcium levels were partially restored, although non-significantly, when the CRT-KO cells were reconstituted with wild-type CRT. CRT-KO cells reconstituted with CRT_Del52_ and CRT_Del52-KDEL_ showed small but non-significant decreases in ER calcium levels compared to those reconstituted with wild-type CRT (Figure 4D, right panel). Changes to ER calcium levels in response to thapsigargin (Tg), a SERCA inhibitor, were similar in the parental *vs.* CRT-KO cells, and in the CRT-KO cells compared with those reconstituted with different CRT proteins (Figure 4E).

Spectrofluorimetry was used to verify the maintenance of ER calcium signals in the different HEK cell lines. Cells were again transfected with the pCIS-GEM-CEPIA1er probe and spectra were recorded in a buffer containing 2 mM Ca^2+^, following excitation at 390 nm. Representative spectra are shown in Figure 5A. Consistent with the flow cytometric measurements, there were no significant differences in the basal ER calcium levels between CRT-KO cells and those reconstituted with wild-type CRT or CRT_Del52_ (Figure 5B). There were also similar changes to ER calcium levels in response to Tg treatment in the CRT-KO cells compared to cells reconstituted with either wild-type CRT or CRT_Del52_ (Figure 5C).

**Figure 5:**
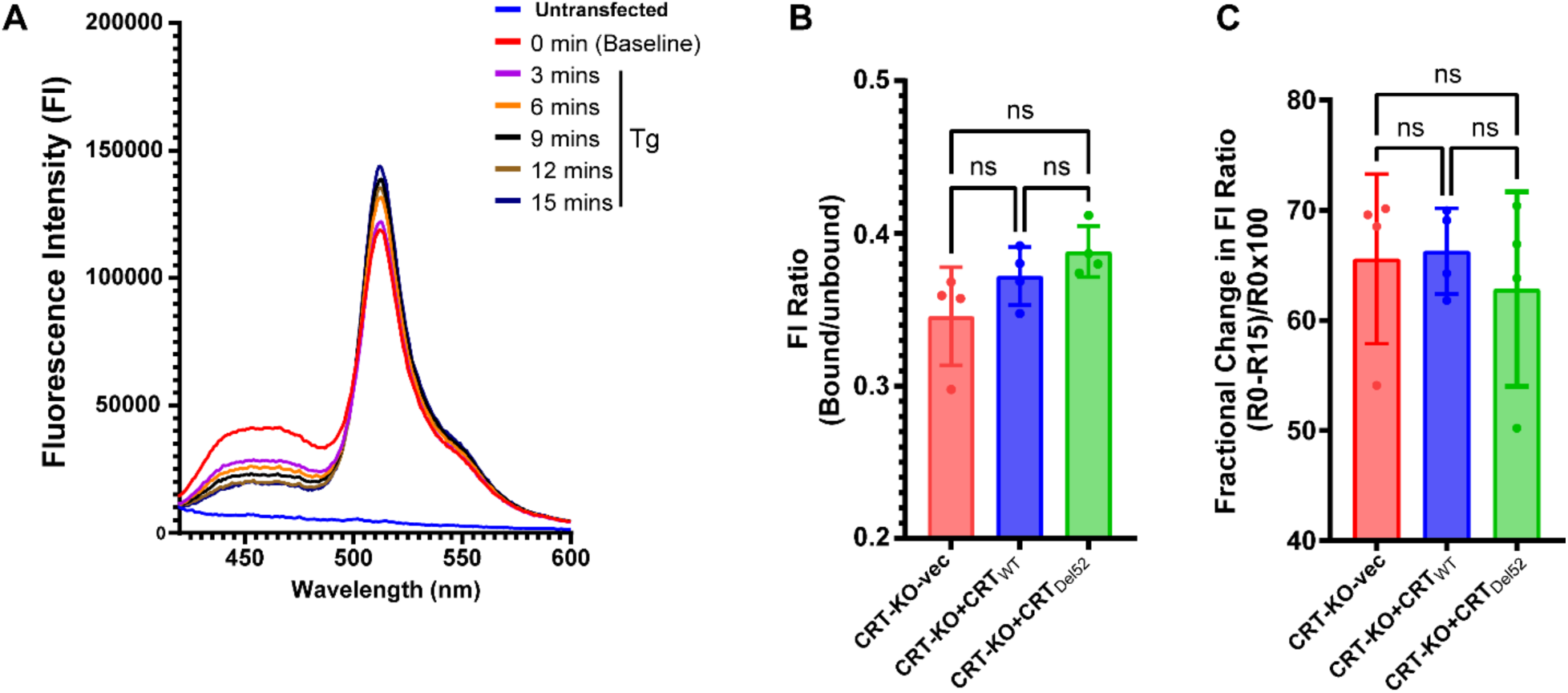
Fluorimetry-based ER calcium measurements indicate similar ER calcium levels in CRT-KO HEK293T cells and those reconstituted with wild-type CRT and CRT_Del52_. HEK cells transiently transfected with a vector encoding the ratiometric GEM-CEPIA1er probe for 72 hours were used to assess ER calcium levels in the presence of 2 mM extracellular Ca^2+.^ Fluorescence emission spectra were recorded following excitation at 390 nm. **A)** Representative emission spectra show changes to Ca^2+^-bound (∼460 nm) and unbound (∼510 nm) peaks at different time points following Tg treatment of cells. The spectrum in blue was obtained with untransfected HEK cells. **B)** Basal Ca^2+^-bound (∼460 nm)/unbound (∼510 nm) fluorescence intensity (FI) ratios were calculated for the indicated HEK cell lines before Tg addition. **C)** The fractional changes to the Bound (∼460 nm)/Unbound (∼510 nm) florescence intensity ratios at 15 mins relative to baseline were calculated as (R_0_-R_15_/R_0_)x100. Statistical significance was determined by repeated measures one-way ANOVA analyses with Tukey’s test using Graph Prism. Data shown are based on 4 independent experiments, indicated by the individual data points within the graphs. ns, not significant

### Similar cytosolic calcium and SOCE signals in a CRT-KO megakaryocyte cell line reconstituted with wild-type CRT or CRT_Del52_

MPN CRT mutations cause thrombopoietin-independent differentiation of megakaryocytes to platelets via the thrombopoietin receptor (MPL) (Araki et al., 2016, Chachoua et al., 2016, Marty et al., 2016, Li et al., 2018, Masubuchi et al., 2020). We, therefore, used a megakaryoblast cell line, MEG-01, to assess the role of CRT deficiency and the CRT_Del52_ mutant on basal cytosolic calcium levels and SERCA inhibition-induced calcium influx into the cytosol. As with the HEK cells (Figure 4), we created a stable CRT knockout clone of MEG-01 cells by CRISPR/Cas9 editing for cytosolic calcium measurements. The cells were subsequently reconstituted with wild-type CRT or CRT_Del52_. Representative immunoblots are shown to demonstrate the successful knockout and reconstitution (Figure 6A). Wild-type CRT reconstitution in MEG-01 CRT-KO cells resulted in expression levels higher than those in the parental cells (Figure 6A, lanes 7 and 8 compared with lanes 1 and 2). In the presence of 2 mM Ca^2+^ in the media, we observed no significant differences in the basal cytosolic calcium levels or CPA-induced calcium release from ER in CRT-KO cells compared to the parental MEG-01 cells (Figure 6C and 6D). Additionally, no significant differences in basal cytosolic calcium were observed in CRT-KO cells compared to those reconstituted with wild-type CRT or CRT_Del52_ (Figure 6E). Furthermore, cytosolic calcium levels in CRT-KO and the two reconstituted cell lines were not statistically different following CPA treatments.

**Figure 6:**
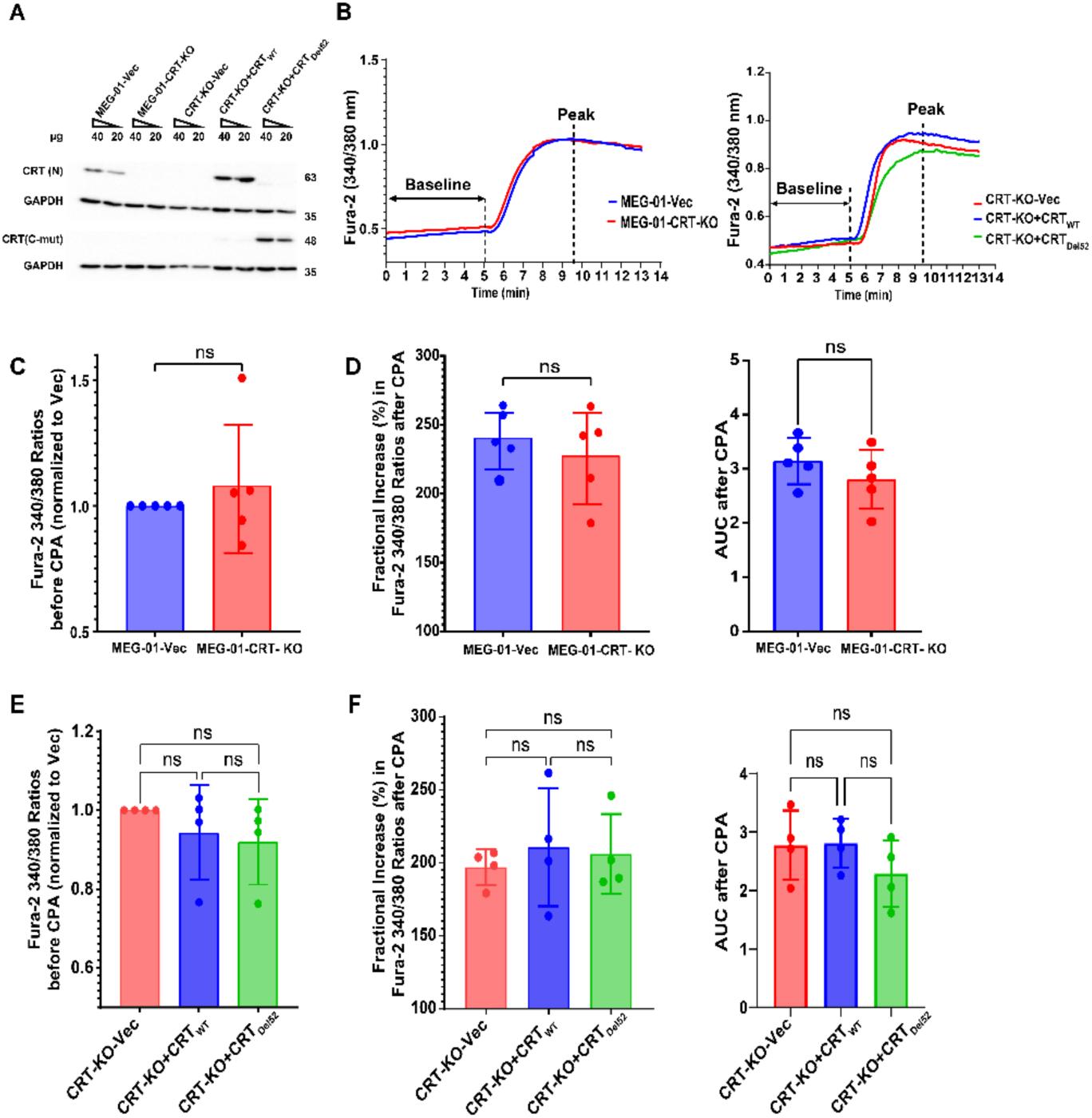
Calcium imaging studies indicate similar cytosolic calcium levels in the megakaryoblastic (MEG-01)cells with CRT-KO and those reconstituted with wild-type CRT or CRT_Del52_. **A)** CRISPR-Cas9 editing of MEG-01 cells using lentiviral constructs encoding calreticulin-targeting gRNA (CRT-KO) or empty vector (Vec). CRT-KO MEG-01 cells were subsequently reconstituted with viral constructs encoding wild-type human CRT or CRT_Del52_ or transduced with empty vector (Vec) as indicated in the Figure panels. Representative immunoblots of CRT expression in the indicated cells detected by the anti-CRT(N) and anti-CRT(C-mut) antibodies. **B-F)** Cells in 2 mM extracellular Ca^2+^ were loaded with the Fura-2/AM dye. Following excitation at 340 or 380 nm and recordings of emission at 510 nm, the emission ratios (340/380) are calculated. **B)** Representative calcium imaging data from indicated cells in the presence of 2 mM extracellular Ca^2+^ and following ER calcium depletion with SERCA inhibitor, cyclopiazonic acid (CPA). **C and E)** Basal cytosolic calcium levels are plotted as the mean 340/380 ratios. Data were normalized relative to the Vec signals. **D and F)** Mean fractional changes to cytosolic calcium levels calculated as (R_peak_-R_baseline_/R_baseline_)x100 and total calcium release determined as the area under the curve (AUC), are plotted. Data shown in C-F are based on 4 or 5 independent experiments with measurements from averaged 50-100 cells per experiment. Statistical analysis was performed using paired t-tests (C and D) or repeated measures one-way ANOVA analyses with Tukey’s test (E and F). Statistical significance was determined based on p-value ≤ 0.05 using GraphPad Prism. ns, not significant.

*CALR* mutations are generally found in BCR-ABL1-negative MPN (Klampfl et al., 2013, Nangalia et al., 2013), whereas MEG-01 cells have the BCR-ABL1 translocation (Ogura et al., 1985), which induces constitutive tyrosine kinase activity. This could influence intracellular calcium homeostasis (Cabanas et al., 2018). To eliminate possible effects of BCR-ABL1 expression on the measured responses (Figure 6), cytosolic calcium signals in the various MEG-01 cell lines were compared with and without treatment of cells with the tyrosine kinase inhibitor, Imatinib. We initially verified the effect of Imatinib on MEG-01 cells by assessing the expression of STAT5 and the formation of phospho-STAT5 using immunoblots. These analyses showed significantly reduced or abrogated phospho-STAT5 expression upon Imatinib treatment (Figure 6-figure supplement 1). Similar to the findings in the absence of Imatinib (Figures 6 and Figure 6-figure supplement 2A, left panel), there were no significant differences in the basal cytosolic calcium levels in Imatinib-treated CRT-KO cells compared to those reconstituted with wild-type CRT and CRT_Del52_ (Figure 6-figure supplement 2B, left panel). Likewise, CPA-induced calcium release from ER was generally similar in CRT-KO cells compared to cells reconstituted with wild-type CRT or CRT_Del52_ both in imatinib-treated and untreated cells (Figure 6-figure supplements 2A and 2B, middle and right panels).

SOCE is an important process for restoring ER calcium and maintaining intracellular calcium levels. In contrast to cytosolic calcium signals in the presence of 2 mM Ca^2+^ in the media, in calcium-free saline, CRT-KO cells exhibited a small but significant increase in basal cytosolic calcium levels relative to the parental cells (Figure 7A, left panel and Figure 7B). Upon ER calcium depletion with CPA, the mean fractional increase in the cytosolic calcium measured in the Ca^2+^-free saline did not differ significantly in parental cells *vs*. CRT-KO cells. The areas under the curve were also not different for the two cell lines (Figures 7A and 7C). However, consistent with the higher basal cytosolic calcium measured in CRT-KO cells in calcium-free saline, the measured SOCE signal (following the addition of 2 mM Ca^2+^ and CPA) was significantly reduced in CRT-KO cells compared to parental cells (Figure 7D). Reconstitution of the CRT-KO cells with either wild-type CRT or CRT_Del52_ tended to reduce basal cytosolic calcium and induce SOCE (Figure 7A, right panel, 7E and 7G), but both trends were non-significant compared to CRT-KO cells. Importantly, there were no significant differences between the wild-type CRT and CRT_Del52_ reconstituted cells, in either their basal or CPA-induced cytosolic calcium signals or in SOCE (Figures 7E-G). In calcium-free saline, cytosolic calcium signals and SOCE induction were generally similar in CRT-KO cells as in the wild-type CRT and CRT_Del52_ reconstituted cells, both in the presence and absence of imatinib (Figure 7-figure supplement 1).

**Figure 7:**
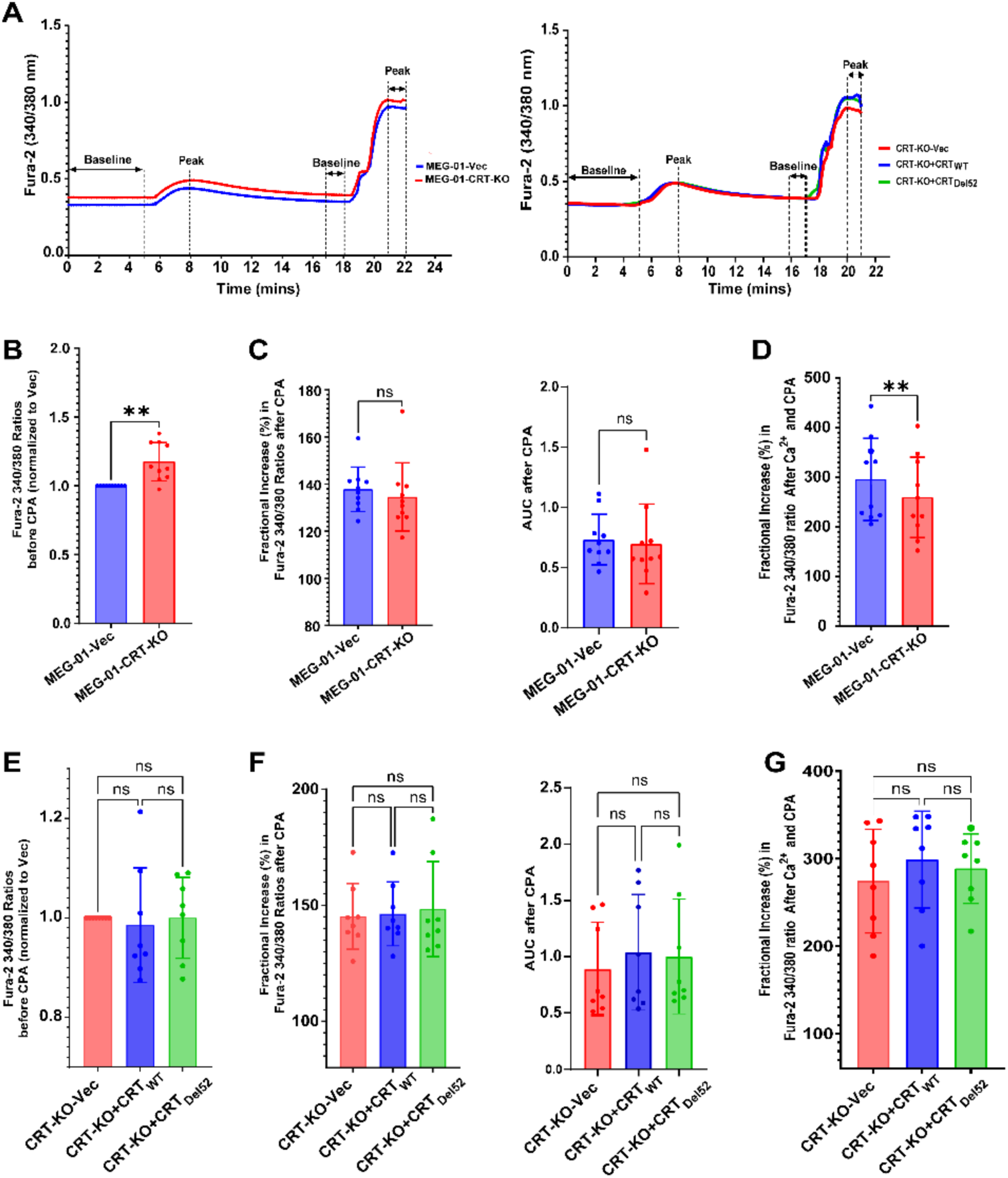
Calcium imaging studies indicate similar cytosolic calcium levels and SOCE in MEG-01 cells with CRT-KO and those reconstituted with wild-type CRT or CRT_Del52_. Cells were loaded with Fura-2/AM dye. Following excitation at 340 or 380 nm and recordings of emission at 510 nm, the emission ratios (340/380) were calculated. **A)** Representative calcium imaging data from cells in the absence of extracellular calcium, followed by the addition of CPA, and subsequent addition of extracellular calcium and CPA as indicated. **B and E)** Basal cytosolic calcium levels are plotted as the mean 340/380 ratios prior to CPA addition. Data are normalized relative to the signals of cells transduced with empty vector (Vec). **C and F)** Fractional changes to cytosolic calcium calculated as (R_peak_-R_baseline_/ R_baseline_)x100 (left panel) and total calcium release measured as the area under the curve (AUC) (right panel) are plotted for the indicated cells, following ER calcium depletion with CPA. **D and G)** Fractional changes to cytosolic calcium following CPA and extracellular calcium additions (2 mM) are calculated as (R_peak_-R_baseline_/ R_baseline_)x100, where R_baseline_ and R_peak_ were measured over 1-minute intervals. Data shown are based on 8-10 independent experiments, and an average of over 20-100 cells in each experiment. Comparisons were performed by paired t-test (panels B-D) and one-way ANOVA analysis (panels E-G) using GraphPad Prism. Statistical significance is based on a p-value ≤ 0.05. ns, not significant.

### CRT deficiency induced numerous transcriptional changes to genes within the calcium signaling pathway

As noted in Figures 4 and 7, some of the measured differences between CRT-KO and parental cells are not fully reversed by reconstitution with wild-type CRT, although the trends are maintained. We predicted that there may be other indirect effects of CRT-KO and reconstitution that affect cellular calcium levels and calcium signaling that could explain some of these measured differences. As proof of concept, we undertook bulk RNA-sequencing to determine differential gene expression in MEG-01 parental *vs.* CRT-KO cells. There were 3078 upregulated and 1790 downregulated genes. The volcano plot and heat map show identified genes that were differentially expressed in CRT-KO MEG-01 compared to parental cells (Figure 8A). Based on pathway analysis, numerous genes involved in cellular calcium signaling were identified. Of specific note is the upregulation of GRP94 encoded by the *HSP90B1* gene, BiP encoded by *HSPA5* gene, and PDIA6 encoded by *PDIA6* gene, which are known to have low affinity calcium binding sites (Lievremont et al., 1997, Macer and Koch, 1988, Biswas et al., 2007, Marzec et al., 2012, Okumura et al., 2021) (Figures 8A and 8B). Other studies have reported similar upregulation of these genes in CRT-KO (Tang et al., 2025) as well as in heterozygous CRT_Del52_ knock-in cells (Figure 8C) (Fosselteder et al., 2023). Upregulation of these proteins may serve a compensatory role in buffering calcium in CRT-KO cells. Additionally, PDIA3, which is shown to regulate the activities of stromal interaction molecule 1 (STIM1) (Prins et al., 2011) and Sarco/endoplasmic reticulum Ca^2+^ ATPase (SERCA2b) (Li and Camacho, 2004), was consistently upregulated in all the studies.

**Figure 8:**
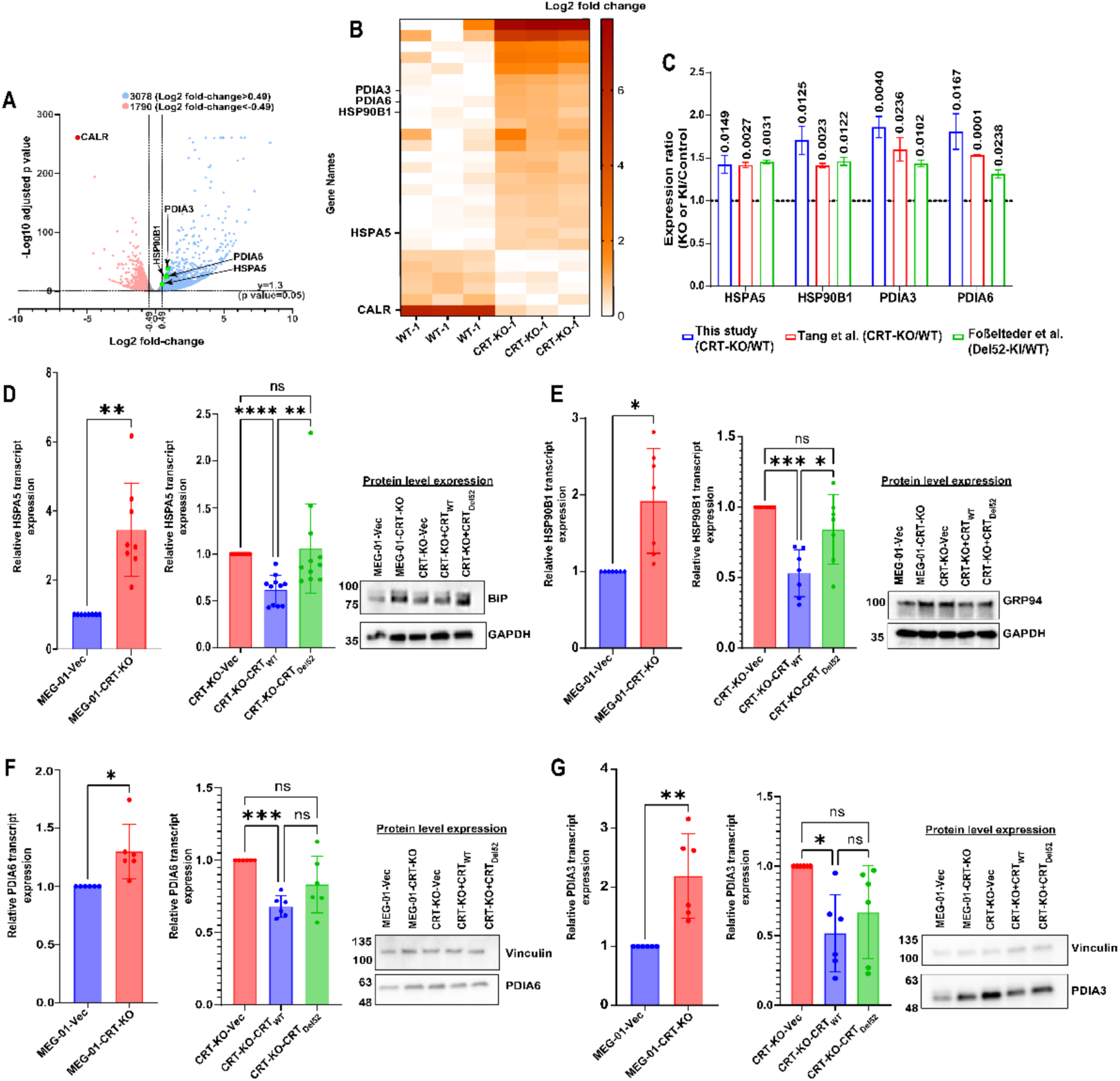
Transcriptional changes to the expression of genes encoding ER calcium binding/regulating proteins in CRT-KO MEG-01 cells. **A)** RNA sequencing analysis was used to identify differentially expressed genes in Meg-01 CRT-KO cells in comparison to parental MEG-01 cells. Volcano plot shows significantly upregulated (blue) and downregulated genes (red) genes identified by RNA sequencing. Genes related to ER calcium binding and homeostasis have been highlighted and labeled. The differentially expressed genes were selected as those with a log2 fold-change of <-0.49 or >0.49 and p values <0.05. **B)** Heat map shows the differential expression of genes encoding ER resident calcium binding proteins or ER calcium regulators that were identified by RNA sequencing in CRT-KO Meg-01 cells compared to parental cells (n=3 biological replicates). **C)** Comparison of expression ratio of *HSPA5, HSP90B1, PDIA3 and PDIA6* in CRT-KO MEG-01 cells (this study) or murine lung cancer cells (Tang et al., 2025), heterozygous CRT_Del52_ knock-in (Del52-KI) cells (Fosselteder et al., 2023) normalized to their expression in respective control cell lines expressing wild type CRT. The normalized expression of various genes identified in each dataset was calculated as a ratio of expression in CRT-KO, CRT_Del52_ knock-in or CRT_Del52_ cells relative to the expression in wild-type CRT-expressing cells. Multiple unpaired t-tests were used to determine statistical significance with the Benjamini-Hochberg FDR correction (p-adjusted) <0.05. **D-G)** Left and middle panels: Relative expression of mRNA transcripts of *HSPA5* (n=8-11), *HSP90B1* (n=7), *PDIA6* (n=6) and *PDIA3* (n=6) genes in the indicated MEG-01 cells determined by RT-qPCR using specific gene primers. Relative gene expression was calculated by normalizing the target gene’s cycle threshold (Ct) to a single or multiple endogenous controls (ACTB, GAPDH and HPRT1) which was used for comparison across different cell lines as indicated. Data show 6 - 11 independent runs with duplicate measurements for each run. Student t-test was used to determine the statistical significance between two groups while multiple group comparison was performed by one-way ANOVA analysis using GraphPad Prism. Statistical significance is based on a p-value ≤ 0.05. ns, not significant. Right panels: Representative immunoblots of BiP, GRP94, PDIA6 and PDIA3 expression in the indicated cells detected by specific antibodies.

We further used real-time PCR with target gene primers (Supplementary Table 2) to validate the expression of the key ER resident proteins identified from the RNA-sequencing data which have roles in calcium binding and regulation of ER calcium homeostasis. Consistent with the RNA-seq data, the expression of *HSPA5, HSP90B1, PDIA6* and *PDIA3* mRNA transcripts was significantly increased in CRT-KO MEG-01 compared to parental cells. Reconstitution of CRT-KO cells with wild-type CRT but not CRT_Del52_ showed a significant decrease in the expression of *HSPA5, HSP90B1, PDIA6 and PDIA3* (Figures 8D-G). Protein levels of BiP, GRP94, PDIA6 and PDIA3 were also assessed in the indicated cells by immunoblots using specific antibodies. Representative immunoblots show increased expression of BiP (Figure 8D, right panel), GRP94 (Figure 8E, right panel), PDIA6 (Figure 8F, right panel) and PDIA3 (Figure 8G, right panel) in CRT-KO cells compared to parental MEG-01 cells as well as increased expression in CRT-KO-Vec cells compared to CRT-KO cells reconstituted with wild type CRT. The effects of CRT_Del52_ in reducing expression were generally less robust than wild type CRT.

## Discussion

Calreticulin is a ubiquitously expressed ER luminal calcium-binding chaperone, with a role in ER calcium storage (Venkatesan et al., 2021b, Michalak, 2024). A number of prior studies have examined the effects of CRT deficiency and over-expression upon cytosolic and ER calcium signals and SOCE (Arnaudeau et al., 2002, Bastianutto et al., 1995, Mery et al., 1996, Nakamura et al., 2001). CRT deficiency in mouse fibroblasts did not affect free ER calcium levels relative to wild type cells (Nakamura et al., 2001). Previous studies using dye-based assays reported impaired calcium binding by the MPN-associated CRT_Del52_ mutant (Ibarra et al., 2022). Yet direct and more quantitative analyses of its calcium-binding properties have been lacking. In this study, we investigated low affinity calcium binding by wild-type human CRT, CRT_Del52_, and the truncation mutants CRT_(1-351)_ and CRT_(1-339)_ using ITC (Figure 3 and Table 1). Our studies show that wild-type CRT, CRT_Del52,_ and CRT_(1-351)_ retain low affinity calcium binding sites, exhibiting average K_D_ values of 574.24, 938.82, and 444.34 µM, respectively. On the other hand, low affinity calcium binding by CRT_(1-339)_ could not be reliably fit. This is the first quantitative characterization of the low affinity calcium binding profiles of wild-type human CRT, CRT_Del52_, and functionally informative truncation mutants.

Although CRT_Del52_ retains at least partial low affinity calcium binding capability, loss of its KDEL retention sequence (Figure 1) could still disrupt ER calcium storage and signaling. Interestingly, the cellular assays reveal that ER and cytosolic calcium levels, as well as SOCE, are similar between CRT-KO cells and those reconstituted with either wild-type CRT or CRT_Del52_ (Figures 4-7). One study has reported decreased ER calcium levels in a human osteosarcoma cell line transiently transfected to over-express a CRT type I mutant, compared to those over-expressing wild-type CRT or a type 2 mutant, which was linked to unfolded protein response (UPR) induction in type I mutant-expressing cells (Ibarra et al., 2022). The same study suggested the contributions of the P and C-domains of wild-type CRT in restoring ER calcium levels in cells expressing CRT_Del52_ (Ibarra et al., 2022). The conditions of those experiments differ from the studies described in our study which uses a CRT-KO background with wild type CRT or CRT_Del52_ reconstitutions to simulate homozygous MPN type I CALR mutations. Achieving endogenous expression levels of wild type CRT is a limitation of our study as in many other studies examining the effects of CRT on cellular calcium signaling. Despite this, our findings are consistent with the theoretical prediction that any excess free ER calcium generated in cells lacking CRT or sufficient ER calcium binding sites would rapidly diffuse out of the ER and be pumped out of the cell.

Related to cytosolic calcium levels and SOCE activation, consistent with our findings (Figures 7G and Figure 7-figure supplement 1), a previous study found no differences in cytosolic calcium levels or SOCE induction in myeloid progenitor 32D cells expressing wild-type CRT and CRT_Del52_ both under conditions of cytokine deprivation and thrombopoietin (TPO) stimulation (Bhuria et al., 2024). On the other hand, studies with cultured megakaryocytes from MPN patients with *CALR* mutations have reported cytosolic calcium abnormalities compared to healthy control cells (Di Buduo et al., 2020, Pietra et al., 2016). In one study, cultured megakaryocytes from patients with type I but not type II *CALR* mutations were found to have high cytosolic calcium and SOCE activation in cultured megakaryocytes (Pietra et al., 2016). In another study, a larger number of cultured megakaryocytes from MPN patients with type I or type II *CALR* mutations displayed spontaneous cytosolic calcium spikes and reduced CRT interactions with stromal interaction molecule I (STIM-1) (reflective of SOCE) compared with megakaryocytes from healthy donors or patients with the JAK2V617F mutation (Di Buduo et al., 2020). Thus, there are differences in the findings of the two studies with primary cultured megakaryocytes, the first of which reported increased SOCE activation in cultured megakaryocytes from patients with type I but not type 2 *CALR* mutations, and the second of which reported increased frequency of cytosolic calcium spikes and reduced STIM-1-CRT binding in cultured megakaryocytes from patients with both type I and type 2 *CALR* mutations (Pietra et al., 2016, Di Buduo et al., 2020). In the absence of engineered controls, sample to sample heterogeneities in primary cells make it difficult to attribute any measured differences to direct rather than indirect effect of CRT mutations on cellular calcium signaling.

We found that CRT-KO HEK cells displayed lower ER calcium compared with the parental HEK cells (Figure 4D, left panel) and also that CRT-KO MEG-01 cells displayed increased cytosolic calcium in the absence of extracellular calcium (Figure 7B). However, reconstitutions with wild-type CRT did not reverse these effects (Figure 4D, right panel and Figure 7E). Transcriptomic analysis identified significant alterations in the expression of genes related to calcium storage and signaling between parental and CRT-KO MEG-01 cells (Figure 8). These changes highlight adaptive or compensatory pathways induced by CRT-KO and emphasize that, when comparing wild-type and CRT-KO cell lines for subcellular calcium levels or responses to SERCA inhibitors, the measured effects might not be based solely on the abilities of CRT to maintain ER and cytosolic Ca^2+^ levels via its calcium binding, storage and buffering capacity. Rather, many other compensatory cellular changes could affect subcellular calcium levels and signaling. In this study, various ER proteins with known calcium binding activities were found to be upregulated in CRT-KO cells (Figure 8A and 8B). GRP94 (Macer and Koch, 1988, Biswas et al., 2007, Marzec et al., 2012), BiP (Lievremont et al., 1997) and our unpublished findings), and PDIA6 (Okumura et al., 2021) are each known to contain low affinity calcium binding sites. CRT-KO cells have significantly induced expression of the genes encoding these proteins. Wild type CRT reconstitution into the CRT-KO cells significantly reduced the expression of these ER proteins, whereas CRT_Del52_ reconstitution only partly and non-significantly reduced the expression of *HSP90B1* and *PDIA6* and did not reduce the expression of *HSPA5* (Figures 8D-F). GRP94, BiP and PDIA6 have functions in ER protein folding and quality control (Hendershot et al., 2024, Okumura et al., 2021). Related to the mechanisms of induction of calcium binding proteins, it is known that CRT_Del52_ has impaired chaperone activity towards its substrates compared with wild type CRT (Arshad and Cresswell, 2018, Schurch et al., 2022, Kaur et al., 2025). This could account for some of the measured differences between wild type CRT and CRT_Del52_ in altering the expression of the ER calcium binding proteins that also function as chaperones and folding factors. However, the induction of GRP94, BiP and PDIA6 in knock-in cells with heterozygous expression of wild type CRT and CRT_Del52_ (Fosselteder et al., 2023), where the wild type CRT appears sufficient for protein folding activities (Kaur et al., 2025, Schurch et al., 2022), points to compensatory effects relevant to calcium signaling. Regardless of the precise mechanisms relevant to induction of the calcium binding proteins, their combined expression, despite protein compositional differences between the KO and reconstituted cells, appears sufficient to maintain ER calcium levels (Figure 4D, right panel), calcium release from the ER (Figure 4E, right panel, and 6F) and cytosolic calcium levels (Figure 6E).

PDIA3 (ERp57) forms a complex with CRT and interacts with and regulates the activities of the ubiquitously expressed SERCA2b (Li and Camacho, 2004). Its expression is also induced by the CRT-KO and suppressed by wild type CRT expression but non-significantly by CRT_Del52_ expression. Altered SERCA2b activity could also affect basal ER calcium levels, contributing to similarities between CRT-KO, and wild type CRT or CRT_Del52_ reconstituted cells.

Collectively, our findings reveal that CRT_Del52_ retains some ability to bind calcium at low affinity sites. CRT-KO cells reconstituted with wild-type or CRT_Del52_ have comparable ER and cytosolic calcium levels and store-operated calcium entry (SOCE). CRT-KO induces broad changes in ER calcium binding protein expression as well as in the expression of ERp57. Overall, we can conclude that a number of factors that contribute to the maintenance of ER calcium levels and regulation of calcium transport into the ER are altered in CRT-KO cells. Thus, any disease-related changes in calcium signaling and homeostasis are likely multifactorial, encompassing transcriptional regulation and altered expression of critical mediators of calcium storage, transport and signaling pathways. These studies provide a more nuanced understanding of CRT’s role in cellular calcium signaling and may inform future therapeutic strategies targeting calcium signaling pathways in hematological malignancies.

## Material and Methods

### DNA constructs

#### Bacterial expression vectors

Wild-type human CRT (accession number P27797) and CRT_Del52_ described earlier (Venkatesan et al., 2021a) was expressed as a fusion protein with the B1 domain of streptococcal protein G (GB1) (Huth et al., 1997) and hexa-histidine tag using the pGB1 vector via ligation-independent cloning, as described earlier (Del Cid et al., 2010, DelProposto et al., 2009). Using the pGB1-h*CALR* plasmid as a template, constructs for expressing his-tagged CRT _(1-351)_ and CRT _(1-339)_ truncation mutants were generated using QuikChange Site-Directed Mutagenesis Kit (Agilent Technologies) following the manufacturer’s protocol and using primers indicated in the Supplemental Table 1.

#### CRT knockdown

The sgRNA for CRT knockdown was designed to target the GGCCACAGATGTCGGGACCT sequence found at the boundary of intron 3 and exon 4 of the *CALR* gene (Mohan et al., 2020). The primers for cloning of sgRNA into pLentiCRISPRv2-BLAST (pLB) vector are given in the Supplemental Table 1.

#### pSIP vectors for CRT expression in human cells

The pMSCV constructs used for expression of wild-type CRT and the CRT_Del52_ described earlier (Venkatesan et al., 2021a) were used as templates to subclone wild-type *CALR* and *CALR_Del52_* cDNA sequences into the pSIP-*ZsGreen* (pLVX-SFFV-IRES-Puro) vector (Bashirova et al., 2020) at the EcoRI and NotI sites using primers given in the Supplemental Table1. The cloning of *CALR* genes at the EcoRI and NotI sites within the pSIP-*ZsGreen* vector resulted in the removal of a fragment carrying *ZsGreen* so that the transduced cells did not express ZsGreen.

#### gRNA resistant versions for reconstitution of CALR expression in KO cells

Silent mutations were introduced into the target sequence for sgRNA in the pSIP-*CALR_WT_ and* pSIP-*CALR_Del52_* constructs by site-directed mutagenesis to make them resistant to Cas9-mediated editing. The QuikChange II XL Site-Directed Mutagenesis Kit (Agilent) was used along with primers listed in the Supplementary Table 1. The PCR reaction was set up according to the manufacturer’s protocol, followed by DpnI treatment for 1 h at 37°C to digest the parental DNA strand. The DpnI-treated product was used to transform XL10-Gold ultracompetent cells. Two sets of primers were used to introduce a total of six silent mutations: C1QF and C1QR primers were used in the first round, and C2QF and C2QR primers were used to introduce further mutations into the DNA templates generated from the first round of mutagenesis.

#### Addition of a KDEL sequence into CRT_Del52_

The pSIP-*CALR_Del52-KDEL_* construct containing a C-terminal KDEL motif was generated using sgRNA-resistant pSIP-*CALR_Del52_* construct as a template and primers given in Supplemental Table 1. The amplicons were cloned at the EcoRI and NotI sites in the pSIP vector. The cloning of *CALR* genes at the EcoRI and NotI sites within the pSIP-*ZsGreen* vector resulted in the removal of a fragment carrying *ZsGreen* so that the transduced cells did not express ZsGreen.

### Cell culture, viral transduction and reconstitution of the expression of CRT mutants

Megakaryoblastic leukemia cell line (MEG-01) from ATCC and Human embryonic kidney (HEK) 293T cell lines were cultured and maintained in RPMI 1640 and DMEM media, respectively, supplemented with 10% heat inactivated fetal bovine serum (FBS, Gibco), antibiotic/antimycotic solution containing penicillin, streptomycin and amphotericin (15240062) respectively at 37°C, 5% CO_2_ and 95% humidity. Wild-type CRT was knocked out in MEG-01 cells and HEK293T cells (CRT-KO) using Cas9 and single guide RNA (sgRNA) targeting GGCCACAGATGTCGGGACCT sequence within the *CALR* gene. For CRISPR/Cas9 mediated CRT knockdown, the HEK293T cells were first transfected with the lentiviral construct containing CRT-targeting sgRNA (pLB-CALR sgRNA) or the empty pLB vector along with the psPAX2 and MD2.G vectors using Lipfectamine™ LTX with PLUS™ reagent (Invitrogen) to make lentivirus. The lentivirus-containing media were harvested, filtered with 0.45 µm syringe filters and added onto the target cells (MEG-01 or HEK293T cells) for infection. The MEG-01 cells were infected by spinoculation at 2500 rpm for 2 hours at room temperature in the presence of polybrene (8 µg/mL), followed by replacement of 80% of virus-containing media with fresh complete media.

For HEK293T cells, the harvested viral supernatant was added to HEK 293T cells and incubated for 2 hours at 37°C, after which 80% of the viral suspension was replaced with complete growth media. Virus-containing media was completely replaced with fresh growth media 24 hours after infection, and the drug selection started with 2.5 µg/mL blasticidin 48 hours post-infection. Following drug selection, the CRT-KO cells were plated at a dilution of one cell per well in a 96-well plate to isolate single-cell clones.

The selected single-cell clones of CRT-KO MEG-01 and HEK293T cells were subsequently reconstituted with wild-type human CRT, CRT_Del52_, or CRT_Del52_ with a C-terminal KDEL sequence using the respective sgRNA-resistant pSIP constructs and the same method of transduction explained above. The cells transduced with the empty pSIP vector without *ZsGreen* (Supplemental Table 1) were also generated to serve as control cells. Cells expressing CRT proteins or those containing empty pSIP vector were selected with 2 µg/mL puromycin. Selected cells were expanded and used for ER calcium measurements.

### Protein expression and purification

The B1 domain of streptococcal protein G (GB1) and his-tagged versions of wild-type CRT, CRT_(1-351)_, CRT_(1-339)_ and CRT_Del52_ were cloned into the pGB1 vector (DelProposto et al., 2009). Chemically-competent Origami B(DE3) *E. coli* cells were transformed with the pGB1 constructs of wild type CRT, CRT_(1-351)_, CRT_(1-339)_ and CRT_Del52._ Single colonies were grown overnight at 37°C. The overnight cultures were inoculated into larger (1-2 L) volumes of LB media and grown in a shaking incubator at 220 rpm and 37°C until the OD_600_ of 0.8. Protein expression was then induced with 200 µM IPTG, and the cultures were further incubated at 16°C overnight on shaking. The bacterial cells were harvested, pelleted, and lysed in 50 mL of HEPES buffer containing (20 mM HEPES pH 7.5, 140 mM NaCl, 1 mM CaCl_2_) by sonication. The lysates were clarified by centrifugation at 10,000 rpm, 4°C for 30 minutes. The supernatant was incubated with nickel-NTA resin (Qiagen, 1018244) for 3 hours at 4°C and purified through a gravity-flow column. The unbound proteins and contaminants were removed by washing with a wash buffer containing 20 mM HEPES pH 7.5, 140 mM NaCl, 1 mM CaCl_2_, 10 mM imidazole. The his-tagged GB1-CRT proteins were eluted with buffer containing 20 mM HEPES pH 7.5, 140 mM NaCl, 1 mM CaCl_2_, 350 mM imidazole. The eluted proteins were concentrated and subjected to fast protein liquid chromatography over a Superdex 200 Increase 10/300 GL column at 4°C in 20 mM HEPES pH 7.5, 140 mM NaCl, 1 mM CaCl_2_ as described previously (Del Cid et al., 2010, Wijeyesakere et al., 2011). Monomer fractions, collected between 14 – 20 mL as shown in Figure 2, were pooled, dialyzed and concentrated to 4 mg/mL in 20 mM HEPES, 10 mM NaCl 1 mM CaCl_2_ pH 7.5 using Amicon ultra concentrators with a molecular weight cut-off of 30,000 Dalton (Millipore) and frozen in 10% glycerol for further use.

### Isothermal titration calorimetry

Isothermal titration calorimetry (ITC) measurements were done using the Nano-ITC SV (TA Instruments). Purified GB1-CRT monomers were dialyzed into 20 mM HEPES, 10 mM NaCl pH 7.5, brought to a concentration of 100 µM, and then pre-incubated with 50-100 µM CaCl_2_ in 20 mM HEPES, 10 mM NaCl pH 7.5 for 30 minutes to block the high affinity calcium binding sites. This was followed by 25 sequential 10µL injections of 20 mM HEPES, 10 mM NaCl, 10 mM CaCl_2_ pH 7.5 into the protein solution maintained at 37°C in a total volume of 1.3 mL to final CaCl_2_ concentrations of 1.6-1.7 mM. The heat changes measured for the sequential injections were curve-fitted to calculate the enthalpies, stoichiometries, and the dissociation constants using NanoAnalyze software.

### Cytosolic calcium measurement and store-operated calcium entry (SOCE) induction

MEG-01 cells and their *CALR*-modified versions were plated on glass coverslips pretreated with 0.01% poly-L-lysine to facilitate cell attachment. In separate experiments, cells were treated with 10 µM Imatinib (Sigma), a tyrosine kinase inhibitor for 2 hours prior to cytosolic calcium measurements. Cells were loaded with 2.5 µM Fura-2/AM dye (Ion Biosciences) supplemented with 0.1% pluronic acid in a physiological salt solution (PSS) containing 140 mM NaCl, 2 mM CaCl_2_, 5 mM KCl, 2 mM MgCl_2_, 10 mM HEPES and 11 mM glucose, and incubated for 60 minutes at 37°C. Live single-cell imaging was done at 20X magnification using an Olympus IX71 inverted microscope equipped with a TILL Polychrome V monochromator light source, a Dual-View emission beam splitter, and a cooled Photometrics Quant-EM camera. Cells were transferred to a perfusion chamber that was mounted on the microscope stage, and PSS with or without 2 mM CaCl_2_ was flowed over cells using a gravity-fed perfusion system. Cytosolic calcium measurements were performed by perfusing Fura-2 loaded cells in the chamber with 2 mM Ca^2+^, followed by the addition of 25 µM cyclopiazonic acid (CPA) (Cayman) into the imaging buffer. To determine cytosolic calcium levels after activating SOCE, cells were initially perfused with Ca^2+^-free PSS containing 0.1 mM EGTA, followed by the addition of 25 µM CPA in Ca^2+^-free PSS and subsequent addition of 2 mM Ca^2+^ and 25 µM CPA in PSS (without EGTA) to reestablish calcium influx. The emission signals corresponding to the 340 and 380 nm excitation were recorded, and the signals resulting from 340/380 nm excitation were calculated using the Meta Fluor software. The basal cytosolic calcium was defined as the 340/380 nm signal ratio before CPA treatment (measured over a 4- or 5-minute interval immediately before CPA addition). The fractional increase in cytosolic calcium levels after ER Ca^2+^ depletion with CPA was calculated as the difference between peak signal (single time point) and baseline (averaged over a 4- or 5-minute interval before CPA addition), as a ratio relative to the baseline. The fractional increase in cytosolic calcium after addition of 2 mM extracellular Ca^2+^ and CPA was calculated as the difference between the peak signal following Ca^2+^ and CPA addition (averaged over a 1-minute time interval following the peak signal) and the baseline (the signal averaged over a 0.8-1-minute interval immediately before the Ca^2+^ and CPA addition) relative to the baseline. Total calcium release after ER Ca^2+^ depletion was reported as the total area under the curve (AUC). Fractional increases were calculated as (Peak-Baseline)/Baseline × 100. This was calculated as peak signals after CPA addition relative to the baseline before CPA (measured over a 4- or 5-minute interval immediately before CPA addition).

### ER calcium measurements in HEK293T cells

*CALR*-modified HEK cells as described in methods were transiently transfected with pCIS-GEM-CEPIA1er (Suzuki et al., 2014); Addgene plasmid # 58217) using Fugene HD reagent (Promega). The transfected cells were harvested after 72 hours. Cells were washed twice with FACS buffer (PBS + 2% FBS) supplemented with 2 mM Ca^2+^ and resuspended in the same FACS buffer. The live-dead stain 7-AAD (BD) was added at 1:200 dilution and the cells were analyzed by flow cytometry using the BD LSRFortessa cell analyzer. The signal for the Ca^2+^-bound GEM-CEPIA1er probe was recorded in the Pacific Blue channel (excitation at 405 nm and emission at 452-455 nm) and the signal for the unbound GEM-CEPIA1er probe was recorded in the AmCyan channel (excitation at 405 and emission at 498 nm). The analysis of flow cytometry data was done in the FlowJo software (v10.10.00). The cells were gated based on the forward (FSC-A) and side scatter (SSC-A), followed by gating on the 7AAD-negative (live) cells, and subsequent gating on the cells positive for the expression of GEM-CEPIA1er probe measured in the Pacific Blue and AmCyan channels. Basal signals were initially recorded, followed by the addition of 5 µM thapsigargin (Tg) and subsequent recording of signals for the treated and untreated cells at four different time points between 2 – 10 minutes. The ratio of mean fluorescence intensity (MFI) of the bound to the unbound signal was used for calculations. In additional experiments, ER calcium levels were measured by spectrofluorimetry in HEK cells transfected with pCIS-GEM-CEPIA1er. Fluorescence emission spectra were recorded between 420-600 nm, following excitation at 390 nm. Basal spectra were initially recorded using 3 million cells suspended in phosphate-buffered saline (PBS) containing 2 mM Ca^2+^, followed by treatment with 5 µM Tg, and additional spectra were recorded at 3 min intervals (3 - 15) mins. The ratios of mean fluorescence intensities of the bound (∼ 460 nm) to unbound (∼ 510 nm) peak signals were calculated in the absence of Tg or at 15 mins following Tg addition to compare the basal signals and the changes following Tg addition.

### Immunoblotting

The HEK293T and MEG-01 cells were harvested and lysed in lysis buffer (50 mM Tris pH 7.5, 150 mM NaCl, 1% Triton X-100, 5 mM CaCl_2_) for 30 minutes at 4°C. The lysates were clarified by centrifugation, and the total amount of protein in the lysates was determined using the BCA reagent kit (Thermoscientific). The volume of lysate containing 20-40 μg of protein was denatured by boiling in 1X SDS loading dye for 5 minutes and subjected to SDS-PAGE. Following electrophoresis, the proteins were transferred onto a PVDF membrane. The membranes were blocked in 5% non-fat dry milk solution in TBST (Tris-buffered saline with Tween-20), followed by immunoblotting and detection for protein expression using primary anti-CRT(N) antibody (Cell Signaling Technology (CST) 12238, 1:10,000 dilution) for detection of CRT_WT_ and anti-CRT(C-mut) antibody for detection of CRT mutants (Venkatesan et al., 2021a). Primary anti-STAT5 (94205S, 1:1000), anti-phospho-STAT5 (93559S, 1:1000), anti-GRP78/BiP (11587-1-AP, 1:10,000), anti-GRP94 (14700-1-AP, 1:10,000), anti-ERp57 (15967-1-AP, 1:10,000) and anti-PDIA6 (18233-1-AP, 1:20,000) were used to probe for the expression of BiP, GRP94, PDIA3 and PDIA6 respectively. Anti-GAPDH (CST 2118, 1:20,000) and anti-vinculin (CST 13901, 1:20,000) were used to probe for GAPDH and vinculin as a loading control. Blots were incubated with horseradish peroxidase-conjugated secondary antibodies (Jackson ImmunoResearch) and developed by chemiluminescence (Pierce™ ECL western blotting substrate, 32106). Immunoblot images were captured using the Azure 300 chemi imager (Azure Biosystems).

### RNA-sequencing

The parental and CRT-KO MEG-01 cells were grown in complete media for 12-18 hours before harvesting. Total RNA was isolated using the RNeasy Plus Mini Kit (Qiagen). This was followed by poly-A selection and enrichment of mRNA using NEB Next Ultra Express RNA kit and a NEB Poly A kit (New England Biolabs). Sequencing was done on the NovaSeq at PE150, targeting 30-40 million reads/sample after cDNA library preparation. Data were pre-filtered to remove genes with 0 counts in all samples. Differential gene expression analysis was performed using DESeq2. The differentially expressed genes were selected as those with a log2 fold-change of <-0.49 or >0.49 and p values <0.05. The results from RNA sequencing were further analyzed in the iPathwayGuide to identify significantly altered proteins related to calcium signaling. The volcano plot was plotted in GraphPad Prism as Log2 fold-change (x-axis) *vs.* -Log10 adjusted p-value.

### Real-time polymerase chain reaction

Total RNA was isolated using the RNeasy Plus Mini Kit (Qiagen) followed by synthesis of complementary DNA using the Superscript III First-Strand Synthesis Kit (ThermoFisher Scientific) following the manufacturer’s protocol. Quantitative RT-PCR was performed on the ABI 7500 FAST Real Time PCR machine (ThermoFisher Scientific, USA) using the SYBR Green PCR Master mix (Applied Biosystems, UK) and gene-specific primers (Supplementary Table 2). Measurement was done in technical duplicates. The expression of target genes was quantified using the 2^−ΔΔCt^ method and normalized to the expression of either one or multiple endogenous control genes including beta-actin (ACTB), glyceraldehyde 3-phosphate dehydrogenase (GAPDH) and hypoxanthine phosphoribosyltransferase 1 (HPRT1). The expression of the target gene in MEG-01 CRT-KO and wild type and CRT_Del52_ reconstitution cells was further normalized to the mean expression of parental MEG-01 or CRT-KO-vec. The calculated relative mRNA transcript expression values (RQ) were plotted using GraphPad Prism.

### Statistical analysis

All data were compiled using Microsoft Excel and analyzed using GraphPad Prism 10.4. A Student’s t-test was used to compare mean differences between two groups, while a one-way ANOVA was used to compare mean differences among more than two groups. The threshold for statistical significance was observed at a p-value ≤ 0.05.

## Data availability

All raw data related to this study will be available at www.immport.org under study accession ID SDY3677.

## Acknowledgements

We thank Grace Pagnucco for her contribution to creating the CRISPR/Cas9-engineered HEK293T CRT-KO cells and Arunkumar Venkatesan for the MEG-01 CRT-KO cells. We would like to extend special thanks and appreciation to Chante Liu and Mathewos Biniam Tebeje from the Satin lab for their support with microscopic calcium measurements. We thank Luis Teran-Rodriguez for assistance with data analysis and Dr. Sivaraj Sivaramakrishnan for many helpful suggestions.

We thank the Advanced Genomic Core (ACG) at the University of Michigan for conducting the library preparation and next-generation sequencing, as well as Rebecca Tagett and the Bioinformatics Core of the University of Michigan Medical School for conducting the RNA sequencing data analysis. We thank Dr. Arthur Sherman for many insightful comments. This work was funded by the National Institute of Health grants (R01 AI123957 to MR and R01DK46409 to LSS) and by the University of Michigan Fast Forward Protein Folding Diseases Initiative.

