## Supplemental Table and Figures for "Molecular and Functional Analysis of Calcium Binding by a Cancer-linked Calreticulin Mutant"

### Supplementary data

This document contains supplementary tables and figures.

**Supplemental Table 1:** List of primers used in this study for cloning of the specified DNA constructs or site-directed mutagenesis as mentioned

| **Name of the construct** | **Primers** | **Purpose** | **References** |
|---|---|---|---|
| pGB1-$CALR_{(1\text{-}351)}$ | *Forward*: 5'-CAGGACGAGGAGCAGAGGTAGAGGAGGATGATGAGGACA-3' | Generate CRT (1-351) truncation mutant by site-directed mutagenesis | This study |
| | *Reverse*: 5'-TGTCCTCATCATCCTCCTCTACCTCTGCTCCTCGTCCTG-3' | | |
| pGB1-$CALR_{(1\text{-}339)}$ | *Forward*: 5'-TAACAAAGGCAGCAGAGAAATAAATGAAGGACAAACAGGAC-3' | Generate CRT (1-339) truncation mutant by site-directed mutagenesis | This study |
| | *Reverse*: 5'-GTCCTGTTTGTCCTTCATTTATTTCTCTGCTGCCTTTGTTA-3' | | |
| pLB-*CALR* sgRNA | *Forward* (**BsmBI**): 5'-CACCGGGCCACAGATGTCGGGACCT-3' | Cloning of *CALR*-sgRNA in pLentiCRISPRv2-BLAST (pLB) vector; the underlined bases correspond to the sgRNA sequence | 10.1074/jbc.RA120.014372 |
| | *Reverse* (**BsmBI**): 5-AAACAGGTCCCGACATCTGTGGCCC-3' | | |
| pSIP Vector minus *ZsGreen* | *Forward* (**PmeI**): 5'-AATTCccg**GTTTAAAC**gccG-3' | Replacement of the *ZsGreen* fragment of the pSIP-*ZsGreen* vector with the PmeI sequence | This study |
| | *Reverse* (**PmeI**): 5'-Gggc**CAAATTTG**cggCCTAG-3' | | |
| pSIP-$CALR_{WT}$ | *Forward* (**EcoRI**): 5'-CGAA**GAATTC**GCCGCCACCATGCTGCTATCCG-3' | Cloning of $CALR_{WT}$ cDNA in pSIP vector; the underlined sequence in forward primer is the Kozak sequence | This study |
| | *Reverse* (**NotI**): 5'-GCATTATT**GCGGCCGC**CTACAGCTCGTCCTTGGC-3' | | |
| pSIP-$CALR_{Del52}$ | *Forward* (**EcoRI**): 5'-CGAA**GAATTC**GCCGCCACCATGCTGCTATCCG-3' | Cloning of $CALR_{Del52}$ cDNA in pSIP vector; the underlined sequence in the forward primer is the Kozak sequence | This study |
| | *Reverse* (**NotI**): 5'-GCATTATT**GCGGCCGC**TCAGGCCTCAGTCCAGCC-3' | | |
| pSIP-$CALR_{Del52+KDEL}$ | *Forward* (**EcoRI**): 5'-CGAA**GAATTC**GCCGCCACCATGCTGCTATCCG-3' | Cloning of $CALR_{Del52+KDEL}$ cDNA in pSIP vector; the underlined sequence in forward primer is the Kozak sequence | This study |
| | *Reverse* (**NotI**): 5'-GA**GCGGCCGC**TCATAATTCATCTTTGGCCTCAGTC-3' | | |
| sgRNA-resistant pSIP-$CALR_{WT}$ and pSIP-$CALR_{Del52}$ | C1QF (*Forward*): ATACAACATCATGTTTGGGCCTGATATCTGTGGCCCTGGCACC | Site-directed mutagenesis to make pSIP-$CALR_{WT}$ and pSIP-$CALR_{Del52}$ constructs sgRNA resistant | This study |
| | C1QR (*Reverse*): GGTGCCAGGGCCACAGATATCAGGCCCAAACATGATGTTGTAT | | |
| | C2QF (*Forward*): CAACATCATGTTTGGGCCTGATATATGCGGACCTGGCACCAAGA | | |
| | C2QR (*Reverse*): TCTTGGTGCCAGGTCCGCATATATCAGGCCCAAACATGATGTTG | | |

Supplemental Table 2: List of primers used for amplification of target genes in qPCR

| **Primers** | **Target gene** | **References** |
|---|---|---|
| *Forward*: 5’-GATGAAGCTCTCCCTGGTGG-3’ | *HSPA5* | This study |
| *Reverse*: 5’-TAGGTGGTCCCCAAGTCGAT-3’ | | |
| *Forward*: 5’-GGAGAGTCGTGAAGCAGTTGAG-3’ | *HSP90B1* | This study |
| *Reverse*: 5'-CCACCAAAGCACACGGAGATTC-3' | | |
| *Forward*: 5’-GTCAGCCACTTGAAGAAGCAGG-3’ | *PDIA3* | This study |
| *Reverse*: 5'-TAGGAACTCGGAGTGAGCCTCA-3' | | |
| *Forward*: 5’-TCAGAAAGGCGAGTCTCCTGTG-3’ | *PDIA6* | This study |
| *Reverse*: 5'-CCTCTTGGCAATGTCCTCGTTG-3' | | |

<u>**SUPPLEMENTARY FIGURES**</u>

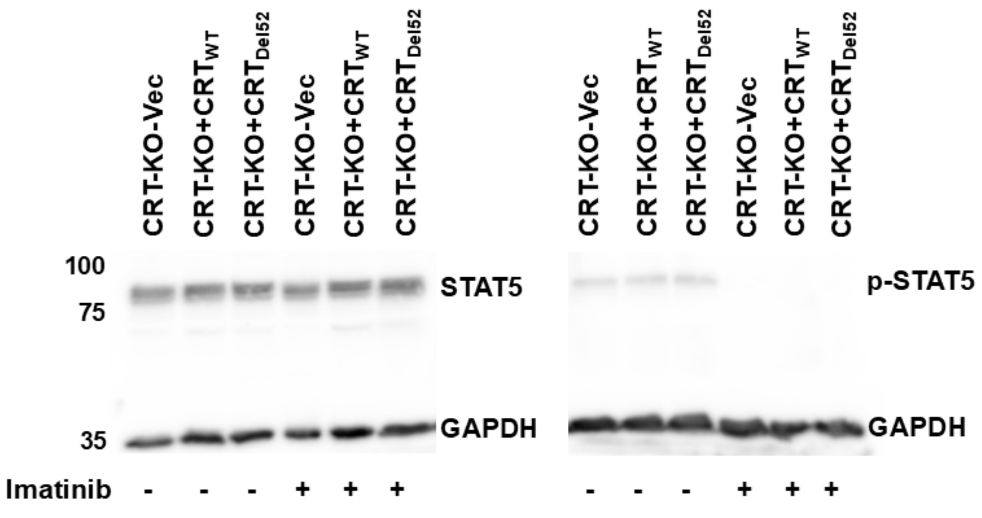


**Figure 6-figure supplement 1: Imatinib treatment reduces p-STAT5 levels.** Representative immunoblots of STAT5 and phospho-STAT5 expression in the lysates of MEG-01 CRT-KO cells reconstituted with wild-type CRT or $CRT_{Del52}$ or transduced with empty vector (Vec), as indicated, with or without imatinib treatment, detected by anti-stat5 and anti-phospho-stat5 antibodies. GAPDH is shown as loading control.

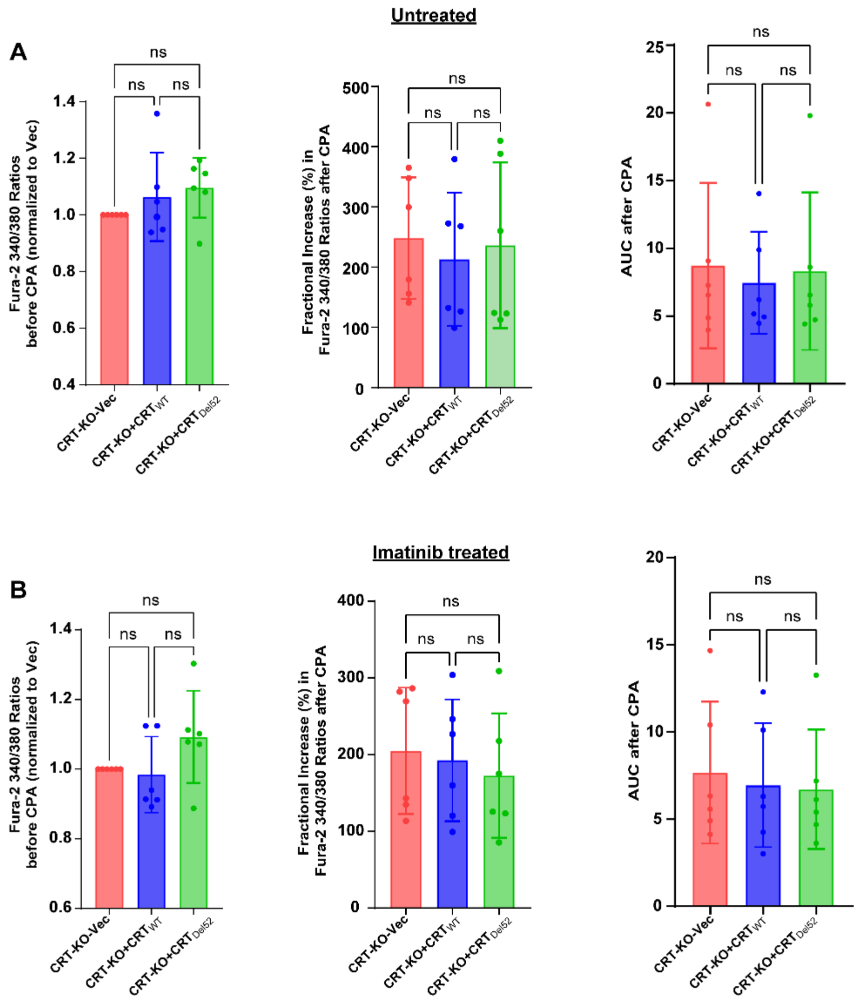


**Figure 6-figure supplement 2: Calcium imaging studies indicate similar cytosolic calcium levels in MEG-01 CRT-KO cells and those reconstituted with wild-type CRT or $CRT_{Del52}$ both with and without imatinib treatment.**

Cells without (**A**) or with imatinib (IM) (**B**) treatment were loaded with Fura-2/AM dye in 2 mM extracellular $Ca^{2+}$ followed by excitation at 340 or 380 nm and recording of emission signals at 510 nm. The basal cytosolic calcium levels before CPA addition are plotted as the mean ratios of emission signals following excitation at 340 or 380 nm (340/380) in cells without imatinib **(A, left panel)** or with imatinib treatment **(B, left panel)**. Mean fractional changes to cytosolic calcium levels calculated as $(R_{peak}-R_{baseline}/R_{baseline})$x100 (Figure 6B) and total calcium release determined as the area under the curve (AUC), following CPA addition to cells without imatinib (**A, middle and right panels, respectively)** or with imatinib **(B, middle and right panels, respectively)** treatment are plotted. Data shown are based on 6 independent measurements, each including 40-70 cells. Repeated measures one-way ANOVA with Tukey's test was used for the statistical analyses. Statistical significance was determined based on p-value ≤ 0.05 using GraphPad Prism. ns, not significant.

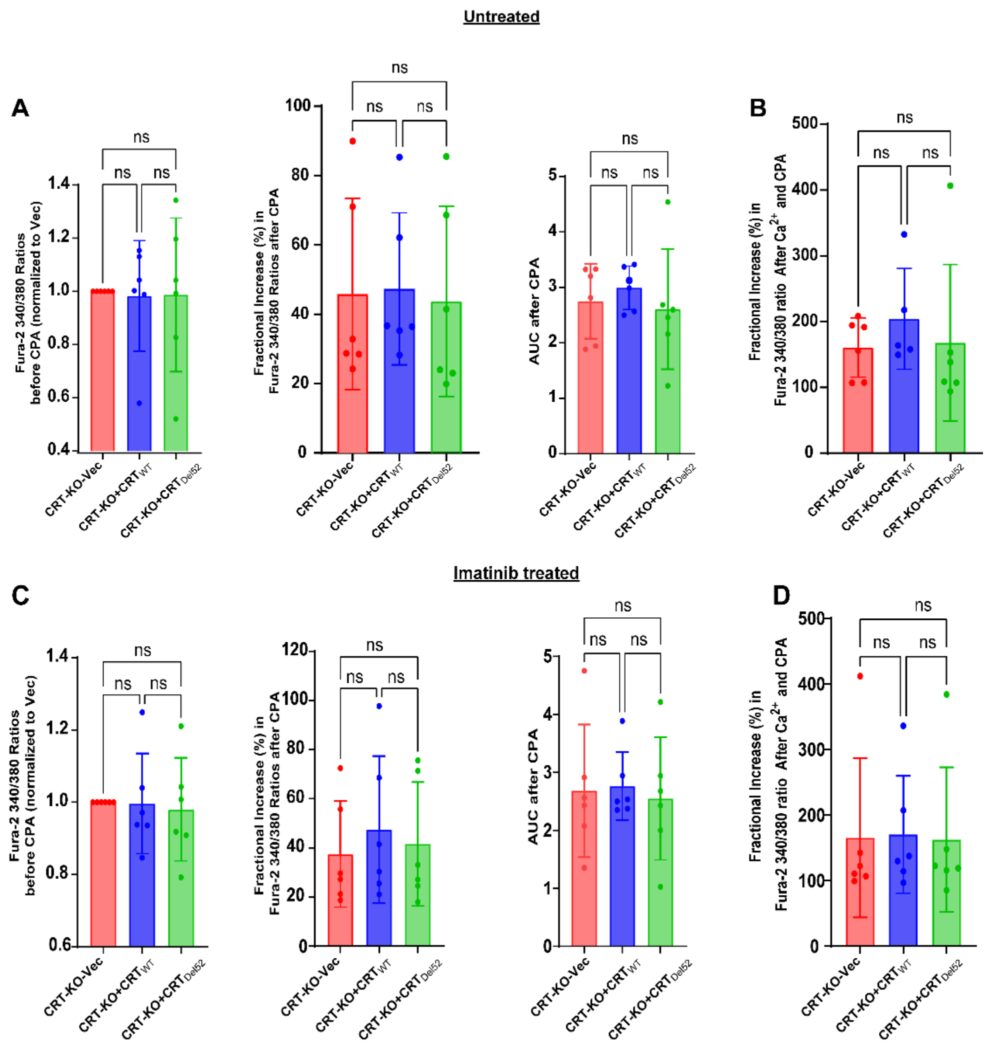


**Figure 7-figure supplement 1: Calcium imaging studies indicate similar cytosolic calcium levels and SOCE in MEG-01 CRT-KO cells and those reconstituted with wild-type CRT or $CRT_{Del52}$ both with and without imatinib treatment.**

Cells without (top panels) or with imatinib (IM) (lower panels) treatment were loaded with Fura-2/AM dye, following excitation at 340 and 380 nm and emission at 510 nm. The ratio of the 340 nm/380 nm signals were calculated. Basal cytosolic calcium levels in the indicated cells without IM **(A, left panel)** or with IM **(C, left panel)** treatment in the absence of extracellular calcium was plotted as the ratios of 340 nm/380 nm signals. Fractional changes to cytosolic calcium and total calcium release measured as the area under the curve (AUC) in the indicated cells following ER calcium depletion with the CPA without IM **(A, middle panels)** or with IM **(C, middle panels**) treatment. **B and D)** Fractional changes to cytosolic calcium following ER calcium depletion with the CPA and subsequent addition of extracellular calcium in the presence of CPA in indicated cells without **(B)** or with IM **(D)** treatment. Data in middle panels of A and C are plotted as a fractional increase relative to baseline following CPA addition. SOCE in the indicated cells is plotted in panels B and D as the fractional increase in signal following the addition of 2 mM extracellular calcium in the presence of CPA. Data shown are based on 6 independent measurements, each with 30-60 cells. Comparisons were performed by one-way ANOVA analysis using GraphPad Prism. Statistical significance is based on a p-value ≤ 0.05. ns, not significant.
